# Optogenetic control of excitatory post-synaptic differentiation through neuroligin-1 tyrosine phosphorylation

**DOI:** 10.1101/788638

**Authors:** Mathieu Letellier, Matthieu Lagardère, Béatrice Tessier, Harald Janovjak, Olivier Thoumine

## Abstract

Neuroligins (Nlgs) are adhesion proteins mediating trans-synaptic contacts in neurons. However, conflicting results around their role in synaptic differentiation arise from the various techniques used to manipulate Nlg expression. Orthogonally to these approaches, we triggered here the phosphorylation of endogenous Nlg1 in CA1 hippocampal neurons using a photoactivatable tyrosine kinase receptor (optoFGFR1). Light stimulation for 24 h selectively increased dendritic spine density and AMPA receptor-mediated EPSCs in wild-type neurons, but not in Nlg1 knock-out neurons or when endogenous Nlg1 was replaced by a non-phosphorylatable mutant (Y782F). Moreover, light stimulation of optoFGFR1 partially occluded LTP. Combined with computer simulations, our data support a model by which Nlg1 tyrosine phosphorylation promotes the assembly of an excitatory post-synaptic scaffold that captures surface AMPA receptors. This optogenetic strategy thus highlights the impact of Nlg1 intracellular signaling in synaptic differentiation and potentiation, while enabling an acute control of these mechanisms.

**Impact Statement:** Orthogonal to the traditional paradigms used to manipulate neuroligin expression level, the optogenetic trigger of tyrosine phosphorylation supports a strong role of endogenous neuroligin-1 in excitatory synaptic differentiation and potentiation.

## Introduction

How early neuronal connections mature into functional synapses is a key question in neurobiology, and adhesion molecules such as neuroligins (Nlgs) are thought to play important roles in this process (Bemben et al., 2015; Craig and Kang, 2007; Südhof, 2008). However, there is an ongoing controversy about the function of Nlgs in synaptic differentiation, arising from divergent results obtained using knock-out (KO), knockdown (KD), and overexpression (OE) approaches. Specifically, whereas Nlg OE or KD bi-directionally affect synapse number, full or conditional Nlg1/2/3 KO does not alter synapse density (Chanda et al., 2017; Chih et al., 2005; Levinson et al., 2005; Prange et al., 2004; Varoqueaux et al., 2006), suggesting that neuroligins are not generally required for synaptogenesis. To address this apparent conflict, experiments that mixed wild type and Nlg1 KO neurons suggested the interesting model that neurons might compete with one another for synapse formation, depending on their intrinsic Nlg1 level (Kwon et al., 2012).

Besides the role of Nlgs in controlling synapse number, there is also a debate about the actual function of Nlgs in regulating basal excitatory synaptic transmission and plasticity. Several studies relying on the expression of Nlg mutants have revealed the potential for Nlg1 to recruit both NMDA receptors (NMDARs) and AMPA receptors (AMPARs) at synapses through extracellular and intracellular interactions, respectively (Budreck et al., 2013; Giannone et al., 2013; Haas et al., 2018; Heine et al., 2008; Letellier et al., 2018; Mondin et al., 2011; Shipman and Nicoll, 2012). However, constitutive or conditional Nlg1/2/3 KO selectively affect basal NMDAR-mediated EPSCs and not AMPARs-EPSCs, and rescue experiments with truncated Nlg1 mutants suggest that the synaptic recruitment of NMDARs requires the intracellular rather than the extracellular domain of Nlg1 (Chanda et al., 2017; Chubykin et al., 2007; Jiang et al., 2017; Wu et al., 2019). Finally, while it is generally accepted that NMDAR-dependent long-term potentiation (LTP) is impaired by Nlg1 KD or KO, the issues of which Nlg1 motifs are important in this process and whether the Nlg1-NMDAR interaction is required, are unclear (Jiang et al., 2017; Kim et al., 2008; Letellier et al., 2018; Shipman and Nicoll, 2012; Wu et al., 2019).

In addition to differences in experimental preparations, these studies relying on the manipulation of the Nlg expression level all have potential biases, including the compensatory expression of proteins in the case of KO (Dang et al., 2018), off-target effects of inhibitory RNAs (Alvarez et al., 2006), and mislocalization of overexpressed Nlgs, e.g. Nlg1 at inhibitory synapses and Nlg2 at excitatory synapses (Chih et al., 2006; Letellier et al., 2018; Nguyen et al., 2016; Tsetsenis et al., 2014). Furthermore, these techniques operate on a long-term basis, i.e. days to weeks, due to slow protein turnover. Hence, there is a pressing need for alternative paradigms allowing for an acute control of Nlg signaling pathways (Jeong et al., 2017) without affecting its expression level. Optogenetics is ideally suited for such purpose and was successfully implemented not only to regulate neuronal excitability and homeostasis, but also for fine tuning protein-protein interactions and signaling pathways in neurons with light (Berlin et al., 2016; Berlin and Isacoff, 2017; Chang et al., 2014; Goold and Nicoll, 2010; Grubb and Burrone, 2010; Mao et al., 2018; Schwechter et al., 2013; Sinnen et al., 2017; Zhang et al., 2011).

To acutely control Nlg1 activity using optogenetics, we aimed at manipulating the phosphotyrosine level of endogenous Nlg1, based on our previous findings that the expression of Nlg1 point mutants in a unique intracellular tyrosine (Y782) affects dendritic spine density and recruitment of PSD-95 and AMPARs (Giannone et al., 2013; Letellier et al., 2018). Here, we show that light stimulation of a photoactivatable receptor tyrosine kinase (optoFGFR1) expressed in hippocampal CA1 neurons increases dendritic spine number, enhances AMPAR-receptor mediated EPSCs, and partially blocks LTP, in a Nlg1 selective fashion, thus demonstrating a major role of the intracellular tyrosine phosphorylation of endogenous Nlg1 in post-synaptic differentiation. Thus, our results show that not only Nlg1 is important for regulating dendritic spine number, but also that the Nlg1 intracellular domain mediates AMPAR recruitment in basal conditions and regulates LTP.

## Results

### Light-stimulation of Nlg1 tyrosine phosphorylation

Using an *in vitro* kinase assay on recombinant GST fused to the intracellular domain of Nlg1, we previously identified several tyrosine kinases able to directly phosphorylate Nlg1, including TRKs and the FGF receptor 1 (FGFR1) (Letellier et al., 2018). To acutely control Nlg1 phosphorylation independently of endogenous, ligand-activated kinases, we thus used here a photoactivatable version of FGFR1 (optoFGFR1) (Grusch et al., 2014) **(Fig. 1A)**. To show that Nlg1 can be acutely phosphorylated by optoFGFR1 in a light-dependent manner, we illuminated COS-7 cells co-expressing recombinant Nlg1 and optoFGFR1 at 470 nm for 15 min using an LED array **(Fig. S1A).** The stimulation of optoFGFR1 by light triggered as much Nlg1 phosphorylation as constitutively active FGFR1 **(Fig. 1B)** (conditions CA and opto+, respectively), indicating potent kinase activation, while samples kept in the dark (conditions light-) did not show significant pTyr levels **(Fig. 1B)**, revealing no unspecific effect of light. Finally, no phosphorylation of the point mutant Nlg1-Y782F was observed upon light application **(Fig. 1B)**, demonstrating that Y782 is the only tyrosine residue on Nlg1 which is phosphorylated by light-gated optoFGFR1.

**Figure 1.**
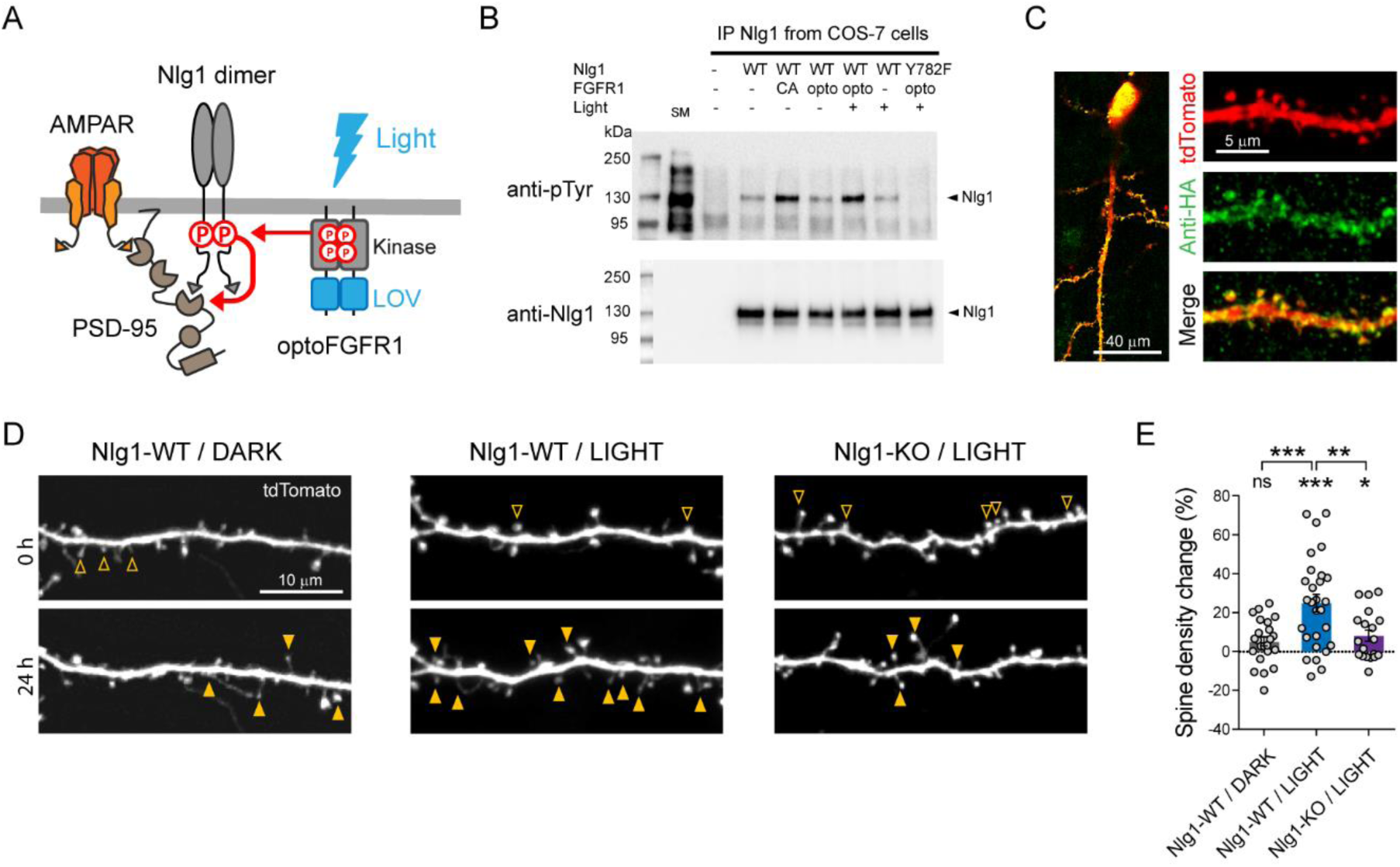
Optogenetic phosphorylation of Nlg1 at residue Y782 increases dendritic spine density. **(A)** Schematic diagram of optogenetically-driven Nlg1 tyrosine phosphorylation using optoFGFR1. Phosphorylated Nlg1 is expected to recruit PSD-95 that serves as a platform for trapping AMPARs. **(B)** Phosphotyrosine immunoblot of proteins extracted from COS-7 cells and immunoprecipitated with anti-Nlg1 antibodies. Cells expressed either no Nlg1, Nlg1 alone, Nlg1 + constitutively active (CA) FGFR1, Nlg1 + optoFGFR1, and Nlg1-Y782F + optoFGFR1. In the first lane, the starting material (SM) from non-transfected cells reveals numerous tyrosine phosphorylated proteins, whereas a single band is present in the Nlg1 IP samples (black arrowhead). Cells were either kept in the dark (-light), or exposed to alternative 470 nm light pulses (1 s light pulse every 1 s) for 15 min (+ light). **(C)** Confocal images of CA1 neurons and dendritic segments showing tdTomato (red) and anti-HA immunostaining (green). **(D)** Confocal images of apical dendrites from electroporated neurons before (0 h) and 24 h after light activation of optoFGFR1. Control slices did not receive light, or received light but were from the Nlg1 KO background. Solid arrowheads point to spines which have appeared, and empty arrowheads to spines which have disappeared in the time interval. **(E)** Normalized spine density for each condition (n = 19-28 dendrites from N = 5-7 cells). Change in spine density was assessed for each condition using paired t-test (\*\*\**P* < 0.001, \**P* < 0.05). Spine density change was compared across conditions using the one-way ANOVA followed by Tukey’s multiple comparison test (\*\*\**P* < 0.001, \*\**P* < 0.01, ns: not significant).

### Light activation of optoFGFR1 increases dendritic spine density

We then examined the impact of triggering Nlg1 tyrosine phosphorylation on synapse morphology and function in organotypic cultures, using confocal microscopy and dual patch-clamp recordings, respectively **(Fig. S1B)**. Using single cell electroporation, we expressed optoFGFR1 with a tdTomato volume marker in CA1 neurons of hippocampal slices obtained from either wild type or Nlg1 KO mice. Immunostained HA-tagged optoFGFR1 was detected throughout dendrites including spines, i.e. at the right location to phosphorylate Nlg1 **(Fig. 1C)**. Dendritic spine density increased by ∼25% in neurons exposed to 470 nm light pulses for 24 h, but remained stable in neurons expressing optoFGFR1 and kept in the dark, or in light-stimulated CA1 neurons from Nlg1 KO slices **(Fig. 1D,E)**, demonstrating that this effect is mediated by light-dependent tyrosine phosphorylation of endogenous Nlg1.

### Light activation of optoFGFR1 enhances basal AMPAR-, but not NMDAR-mediated EPSCs

At the electrophysiological level, neurons expressing optoFGFR1 and exposed to light for 24 h exhibited ∼200% larger AMPAR-mediated EPSCs upon stimulation of Schaffer’s collaterals compared to non-electroporated neighbors that also received light **(Fig. 2A)**, or to neurons expressing optoFGFR1 and kept in the dark **(Fig. 2B,D)**. In contrast, there was no significant effect of optoFGFR1 expression and/or light on NMDAR-mediated EPSCs **(Fig. 2C,E)**. Importantly, the light-induced increase AMPAR-mediated EPSCs was not observed in CA1 neurons from Nlg1 KO slices **(Fig. 2B,D)**, demonstrating that this effect involves the selective tyrosine phosphorylation of Nlg1. The paired-pulse ratio was not changed by optoFGFR1 expression or light exposure, suggesting that presynaptic function was unaltered **(Fig. S2A,B).** Although the critical tyrosine residue belonging to the gephyrin-binding motif is conserved in Nlg2 and Nlg3 (Poulopoulos et al., 2009), where it can also be phosphorylated (Letellier et al., 2018), no effect of optoFGFR1 stimulation was observed on inhibitory currents recorded in CA1 neurons **(Fig. S2C,D)**. Together, these data demonstrate that the phosphorylation mechanism is specific to the Nlg1 isoform at excitatory post-synapses, and selectively affects AMPAR recruitment.

**Figure 2.**
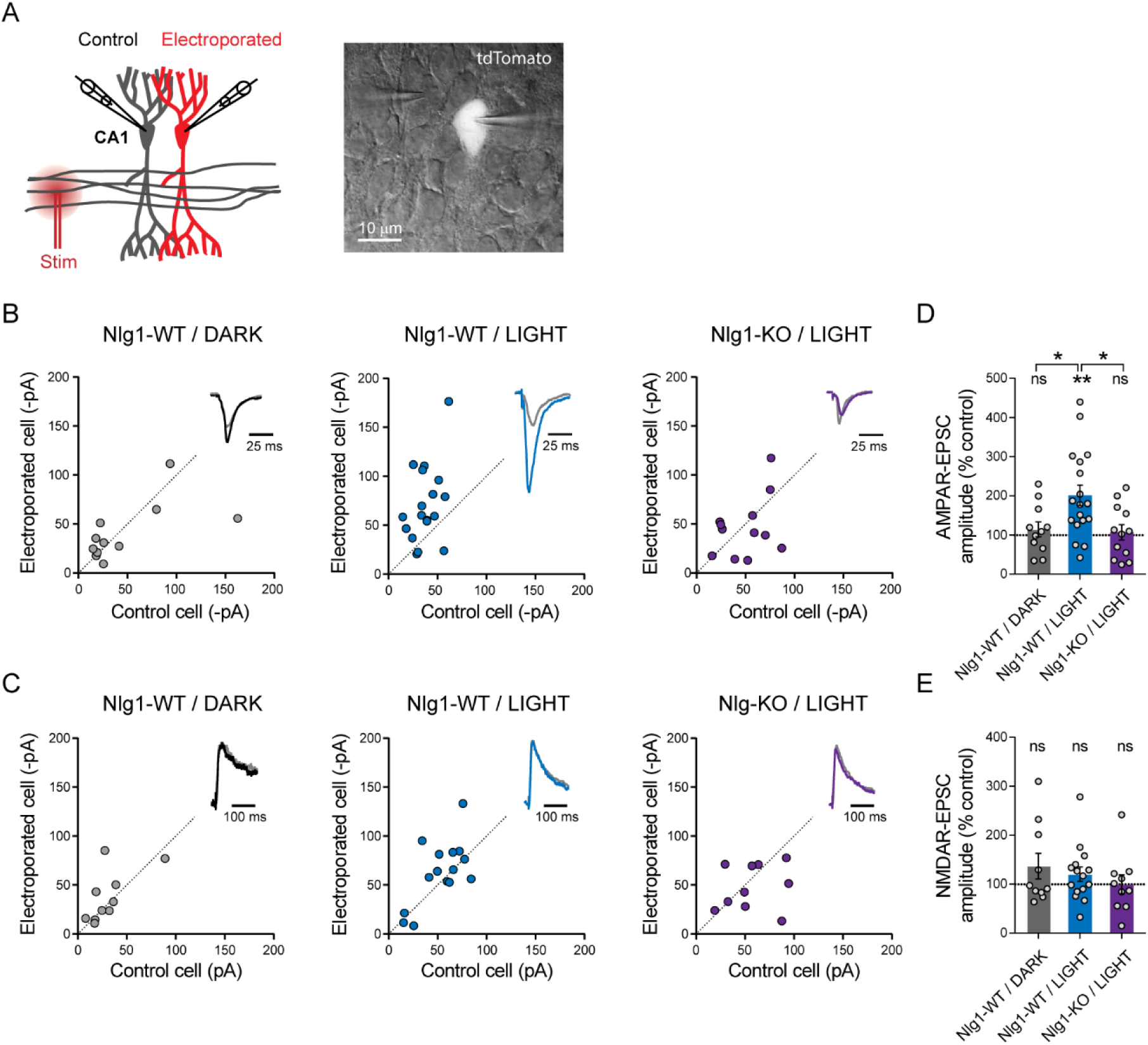
Light activation of optoFGFR1 in CA1 neurons selectively increases AMPAR receptor-mediated EPSCs. **(A)** Dual patch-clamp recordings of AMPAR- and NMDAR-mediated currents upon stimulation of Schaffer’s collaterals were made at holding potentials of -70 and +40 mV (respectively) in electroporated neurons and non-electroporated counterparts. The image shows 2 patched neurons in the CA1 area, the brighter one expressing optoFGFR1 + tdTomato. **(B, C)** Scatter plots of AMPAR- and NMDAR-mediated EPSCs, respectively, in neurons expressing optoFGFR1 compared to paired unelectroporated neurons (control cell), in the indicated conditions. Representative traces (color) normalized to control (grey) are shown as insets. **(D, E)** Average of AMPAR- and NMDAR-mediated EPSCs in the three conditions, normalized to the control (100%). Data were compared to the control condition by the Wilcoxon matched-pairs signed rank test, and between themselves using one-way ANOVA followed by Tukey’s multiple comparison (\*\**P* < 0.01, *P < 0.05, ns: not significant).

### The intracellular Y782 residue is involved in light-induced effects

To verify that optoFGFR1 was specifically phosphorylating the Nlg1 Y782 residue in neurons, we adopted a replacement strategy (Letellier et al., 2018) by co-electroporating optoFGFR1, Nlg1-shRNA, and Nlg1 rescue constructs in slices from WT mice **(Fig. 3A,B)**. This led to basal AMPAR- and NMDAR-mediated EPSCs in the dark matching those measured in paired non-electroporated neurons expressing endogenous Nlg1 levels **(Fig. 3C-F)**. In CA1 neurons expressing rescue Nlg1-WT, light exposure induced again a 25% increase in dendritic spine number **(Fig. S3A,B)**, as well as a 200% increase in AMPA- (but not NMDA-) receptor mediated EPSCs compared to control non-electroporated neurons **(Fig. 3C-F)**. The increase in spine density and AMPAR-mediated EPSCs by optoFGFR1 activation was not observed in CA1 neurons expressing Nlg1-WT and kept in the dark, or in neurons expressing Nlg1-Y782F and exposed to light **(Figs. S3A,B and 3C,E)**. Thus, those effects are likely mediated by phosphorylation of the Nlg1 Y782 residue induced by the photo-activation of optoFGFR1.

**Figure 3.**
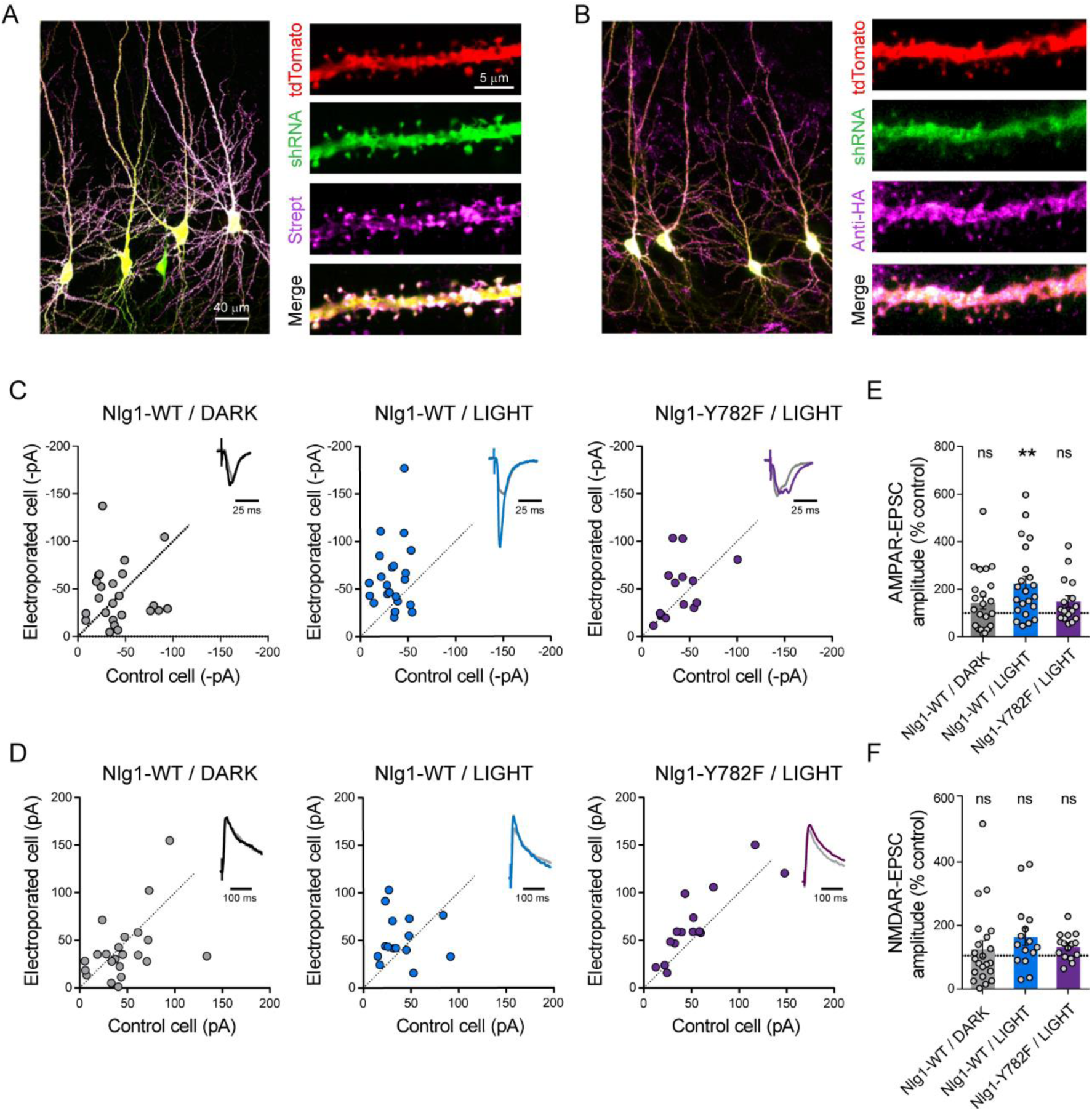
The light-induced increase in AMPA receptor mediated EPSCs is specific to Y782 phosphorylation in Nlg1. CA1 Neurons were co-electroporated with plasmids encoding tdTomato, shRNA against endogenous Nlg1 containing a GFP reporter, shRNA resistant AP-tagged Nlg1-WT or -Y782F, biotin ligase (BirA^ER^), and HA-tagged optoFGFR1. **(A, B)** Confocal images showing tdTomato (red) and GFP (green). Biotinylated Nlg1 and optoFGFR1 were stained in different slices using streptavidin-Atto647 and anti-HA antibody, respectively (magenta). **(C, D)** Scatter plots of AMPAR- and NMDAR-mediated EPSCs, respectively, for electroporated versus paired non-electroporated neurons (control cell) in the indicated conditions. Representative traces (black, blue or violet) normalized to control (grey) are shown as insets. **(E, F)** Average of AMPAR-and NMDAR-mediated EPSCs, respectively, normalized to the control (100%) in the different conditions. Data were compared to the control condition by the Wilcoxon matched-pairs signed rank test (\*\**P* < 0.001).

### Light activation of Nlg1 tyrosine phosphorylation impairs LTP

Finally, we asked whether the increase of basal AMPAR-mediated currents induced by Nlg1 phosphorylation could partially occlude long term potentiation (LTP). CA1 neurons expressing optoFGFR1 and pre-exposed to light showed a ∼2-fold reduction in the LTP plateau level compared to control non-electroporated neighbors **(Fig. 4A,B)**. To quantitatively interpret those results, we carried out computer simulations describing membrane diffusion and synaptic trapping of individual AMPARs, based on a previous framework using realistic kinetic parameters (Czöndör et al., 2012) **(Fig. S4A-C)**. This model is in line with experiments showing that hippocampal LTP primarily involves the capture of extra-synaptic AMPARs (Granger et al., 2013; Penn et al., 2017). We mimicked LTP by introducing a step decrease in the apparent off-rate between AMPARs and the PSD scaffold **(Figs. 4C and S4D)**. The simulations matched very well experimental LTP, both in terms of kinetics and plateau value (∼270%), supporting this diffusion/trap model **(Figs. 4A and S4D)**. To mimic the effect of Nlg1 phosphorylation on postsynaptic density (PSD) assembly and AMPAR recruitment (Letellier et al., 2018), we raised the AMPAR/scaffold binding rate, resulting in a ∼2-fold increase of basal synaptic AMPAR number **(Fig. S4E)** reproducing the experimental data **(Fig. 2B,D)**. In response to the same LTP simulation, the relative increase in AMPAR number now reached only ∼190%, as in optoFGFR1 experiments **(Fig. 4A)**. Thus, the partial occlusion of LTP observed upon optoFGFR1 stimulation can be explained by a high initial recruitment of synaptic AMPARs, which depletes the extra-synaptic AMPAR reservoir necessary for LTP. Accordingly, we previously reported an almost complete occlusion of LTP upon replacement of endogenous Nlg1 by a Y782A mutant which phenocopies maximally phosphorylated Nlg1 and increases AMPAR-mediated EPSCs by ∼4-fold (Giannone et al., 2013; Letellier et al., 2018). Overall, our model predicts a negative correlation between basal synaptic AMPAR number and the ability to respond to LTP **(Fig. S4F)**, that perfectly fits the experiments **(Fig. 4D)**. These data suggest that Nlg1 tyrosine phosphorylation impairs LTP by promoting high initial synaptic AMPARs levels.

**Figure 4.**
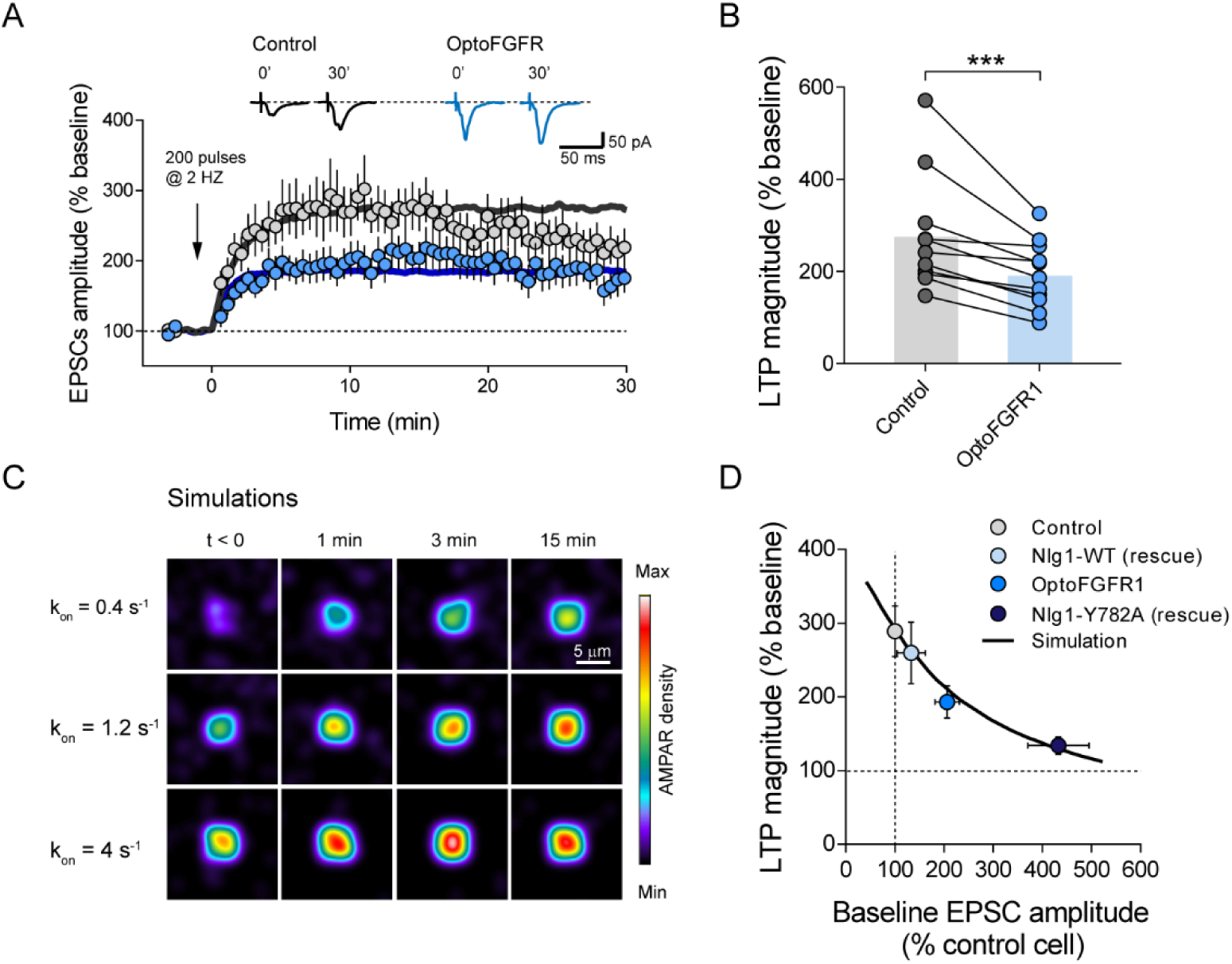
Light activation of Nlg1 phosphorylation by optoFGFR1 reduces LTP. **(A)** Average AMPAR-mediated EPSCs in CA1 neurons expressing optoFGFR1 (blue circles) or in non-electroporated neighbors (grey circles), all pre-exposed to light for 24 h, upon LTP induction at time 0 using a pairing protocol. Sample traces are shown at time 0 and 30 min after LTP induction. The solid lines show averages from 10 computer simulations for each condition. **(B)** Distribution of the long-term plateau of AMPAR-mediated EPSC in the two conditions (6-10 min after LTP induction), expressed as a percentage of the baseline level. Data were compared to the control condition (unelectroporated) by Wilcoxon matched-pairs signed rank test (\*\*\**P* < 0.001). **(C)** Heat maps representing simulations of AMPAR accumulation at a single synapse over time. LTP is mimicked by decreasing the AMPAR/scaffold off-rate at time zero, causing a diffusional trapping of AMPARs from extra-synaptic pools. Nlg1 phosphorylation is mimicked by increasing the initial AMPAR/scaffold on-rate, resulting in higher basal AMPAR level and lower LTP (relatively to baseline). **(D)** Relationship between basal synaptic AMPAR content and LTP plateau level (% of baseline). Experimental points (circles) were obtained from non-electroporated neurons (grey), neurons expressing opoFGFR1 and stimulated with light (blue), or neurons co-expressing shRNA to Nlg1 and either Nlg1-WT (light blue) or Nlg1-Y782A (dark blue) (data taken from (Letellier et al., 2018)). Basal AMPAR-mediated EPSCs were normalized to match a synaptic level of 33 AMPARs in the control condition (Levet et al., 2015). The solid line represents simulated data generated by varying the AMPAR/scaffold binding rate from 0.1-10 s^-1^, while LTP was mimicked by a drop of k_off_ from 0.02 s^-1^ to 0.004 s^-1^.

## Discussion

Orthogonal to the traditional paradigms used to manipulate Nlg expression level or replace Nlg isoforms with truncated or mutated versions, this novel optogenetic approach allows for a fine tuning of the tyrosine phosphorylation of endogenous Nlg1, revealing a strong role of Nlg1 intracellular signaling in excitatory post-synapse differentiation. Our results show that Nlg1 phosphorylation specifically regulates dendritic spine number, mediates AMPAR recruitment in basal conditions, and impairs LTP.

Together, our results support a mechanism by which, in its tyrosine phosphorylated state, Nlg1 preferentially recruits intracellular PDZ domain containing scaffolding proteins including PSD-95 (Giannone et al., 2013; Jeong et al., 2019), associated with a morphological stabilization of dendritic spines (Cane et al., 2014) and serving as slots for the diffusional trapping of surface AMPARs (Czöndör et al., 2013; Mondin et al., 2011). In contrast, NMDAR-mediated EPSCs are not affected by Nlg1 phosphorylation, supporting the concept of a direct extracellular coupling between Nlg1 and GluN1 (Budreck et al., 2013; Shipman and Nicoll, 2012). The selective effects of endogenous Nlg1 tyrosine phosphorylation on dendritic spine density, AMPAR-receptor mediated EPSCs, and LTP are in close agreement with our previous observations based on the KD + rescue of Nlg1 point mutants, in particular the Nlg1 Y782A mutant, which promotes synaptic recruitment of PSD-95, strongly enhances basal AMPAR-mediated EPSCs, and totally blocks LTP through synapse unsilencing mechanisms (Letellier et al., 2018). Importantly, optogenetic Nlg1 phosphorylation induces a similar response as PSD-95 overexpression by increasing spine density, enhancing AMPAR-but not NMDAR-dependent transmission, and occluding LTP (Ehrlich and Malinow, 2004; El-husseini et al., 2000; Stein et al., 2003), which further supports our model.

Our computer simulations of AMPAR diffusional trapping at PSDs provide a simple framework to interpret the experimental data. By triggering PSD scaffold assembly, Nlg1 tyrosine phosphorylation provides synapses with fresh surface-diffusing AMPARs. These already potentiated (or unsilenced) synapses are thus less prone to respond to the LTP stimulation, because extra-synaptic pools of AMPARs have been consequently depleted (Granger et al., 2013; Penn et al., 2017). A strong role of Nlg1 / PSD-95 interaction in synaptic function is also supported by a recent study showing that PKA-mediated phosphorylation of the S839 residue located near the C-terminal PDZ domain binding motif dynamically regulates PSD-95 binding, affecting both dendritic spine number and AMPAR-mediated miniature EPSCs (Jeong et al., 2019).

Although the tyrosine residue in Nlg1 belongs to a gephyrin-binding motif which is highly conserved among the other Nlg isoforms and controls gephyrin binding (Giannone et al., 2013; Poulopoulos et al., 2009), optoFGFR1 stimulation did not affect evoked inhibitory currents, suggesting that the phosphorylation of Nlg2 or Nlg3 either did not occur, or did not modify the recruitment of the gephyrin scaffold and associated GABA_A_ receptors. The lack of effects of optoFGFR1 stimulation in neurons from Nlg1 KO slices confirms that it is indeed Nlg1 phosphorylation which is causing the observed increases in dendritic spine density and AMPAR-mediated EPSCs. It might be interesting to apply similar optogenetic approaches to control the phosphorylation of other intracellular Nlg1 residues including S839 and T709 by engineering photoactivatable versions of PKA and CamKII, respectively (Bemben et al., 2013; Jeong et al., 2019), or inhibitors of those kinases (Murakoshi et al., 2011) expecting to alter Nlg1 trafficking, and thereby synaptic function and potentiation. Other phosphorylation sites have been reported in Nlg2 and Nlg4 which might also be interesting to target with such light-gated kinases (Antonelli et al., 2014; M. a. Bemben et al., 2015).

Because the optoFGFR1 is lacking a ligand-binding domain, its light-activation is expected to by-pass the endogenous regulation of Nlg1 tyrosine phosphorylation, which involves the Trk family of tyrosine kinases (Letellier et al., 2018) that are responding to intrinsic ligands (BDNF and NGF) (Harward et al., 2016). Photoactivatable versions of Trks have been reported and their stimulation with light for 48 hr induces neurite outgrowth in DIV1-3 dissociated neurons and de novo formation of axonal filopodia within 30 min (Chang et al., 2014), but the effects on spine formation and synaptic transmission in mature neurons have not been measured yet. Short-term photoactivation of another tyrosine kinase, EphB2, leads within seconds to the retraction of non-stabilized dendritic filopodia (Mao et al., 2018) and within minutes to the induction of new filopodia by activating actin polymerization (Locke et al., 2017). Those effects are likely to obey different downstream signaling pathways than the ones we report here and which highly depend on Nlg1 and the associated PSD scaffold.

Our data demonstrating the critical role of a single tyrosine residue located in the middle of the intracellular motif are difficult to reconcile with a previous report showing that a Nlg3 construct with a 77-aa intracellular truncation (thus removing the motif containing the tyrosine) can still rescue AMPAR-mediated synaptic transmission upon Nlg1/2/3 KD (Shipman et al., 2011). Moreover, whereas Nlg1 KO was shown to affect primarily basal NMDAR-mediated synaptic transmission, we find instead strong effects of acute Nlg1 tyrosine phosphorylation on basal AMPAR-mediated EPSCs, and no alteration of NMDAR-dependent EPSCs. The fact that AMPAR-mediated EPSCs are not altered in the Nlg1 KO (Chanda et al., 2017) may result from the compensatory expression of scaffolding or adhesion molecules, in particular Nlg3 (Dang et al., 2018), which also interacts with PSD-95. This would explain the fact that a dual Nlg1/3 (and triple Nlg1/2/3) KO are required to alter AMPARs levels and AMPAR-mEPSCs in cultured neurons (Chanda et al., 2017). In contrast, a compensatory expression of Nlg3 which does not interact extracellularly with NMDARs (Budreck et al., 2013; Shipman and Nicoll, 2012) is not expected to rescue the decrease in NMDAR-EPSCs caused by Nlg1 KO.

Finally, while LTP is impaired upon intracellular stimulation of Nlg1 tyrosine phosphorylation, another study finds that LTP can be rescued in acute slices from Nlg1/2/3 cKO upon expression of a GPI-anchored Nlg1 lacking the entire intracellular domain (Wu et al., 2019), and thus the C-terminal PDZ domain binding motif which we find important for anchoring AMPARs through PSD-95 (Letellier et al., 2018; Mondin et al., 2011). While the differences might come from the use of different experimental preparations (acute vs organotypic slices) and perturbation approaches (KD or KO, each with specific timing with respect to the synaptogenesis period), we believe that our approach allowing for an acute control of a signaling mechanism associated with endogenous Nlg1, demonstrates a strong role of the Nlg1 intracellular domain in synaptic function. Besides clarifying the role of neuroligin-1 at excitatory synapses, the optogenetic phosphorylation of neuroligin-1 provides the exciting opportunity to control in time and space synaptic connectivity and function, and has a therefore a great potential for investigating the causality between synaptic plasticity and learning processes as well as the possible contribution of neuroligins to neuropsychiatric behaviors (Bourgeron, 2015).

## Materials and Methods

### Constructs

Plasmids for BirA^ER^ and AP-Nlg1 harboring both extracellular splice inserts A and B were kind gifts from A. Ting (Stanford University, CA). Short hairpin RNA to murine Nlg1 (shNlg1) was a generous gift from P. Scheiffele (Biozentrum, Basel). shRNA-resistant AP-tagged Nlg1 and Nlg1-Y782F were described previously (Chamma et al., 2016; Letellier et al., 2018). FGFR1-Flag was a generous gift from L. Duchesne (Université de Rennes). To generate constitutively active (CA) FGFR1-Flag, the V561M mutation was introduced using the In-Fusion® HD Cloning Kit (Takara Bio) and the following primers: 5’TACTCCATAATGACATAAAGAGG3’ and 5’TGTCATTATGGAGTACGCCTC3’. optoFGFR1 bearing an N-terminal myristoylation motif to attach to the membrane, and a C-terminal HA-tag was described previously(Grusch et al., 2014). In this construct, the extracellular FGF binding domain has been removed, and a light-oxygen voltage sensing (LOV) domain is fused to the C-terminus, such that stimulation with blue light induces dimerization of the FGFR1 intracellular domain and subsequent kinase activation in a ligand-independent manner. The tdTomato plasmid was a generous gift from R. Tsien (UC San Diego, CA).

### COS-7 cell culture, transfection, and light illumination

COS-7 cells were cultured in DMEM (Eurobio) supplemented with 1% glutamax (GIBCO), 1% sodium pyruvate (Sigma-Aldrich), 10% Fetal Bovine Serum (Eurobio). For IP experiments, COS-7 cells were plated in 6-well plates at a density of 70,000 per well. After 1 day, cells were transfected or not with Nlg1 and FGFR1 constructs, and left under a humidified 5% CO_2_ atmosphere (37°C) for 2 days. For cells expressing optoFGFR1, all steps before sample loading were done in the dark. COS-7 cells were exposed to light pulses of 1 s every other second for 15 min, by placing the plates on a custom-made rectangular array comprising 8 × 12 light emitting diodes (LEDs) (470 nm, 15 mW each), powered by a 24 V DC supply, and driven by an internal Arduino® Leonardo pulse generator. The array was covered with a 5-mm thick white Plexiglas sheet to dim the emitted light power by ∼100-fold (2.5 µW/mm^2^).

### Immuno-precipitation, SDS–PAGE, and immunoblotting

Cells were treated with 10 µM pervanadate for 15 min before lysis to preserve phosphate groups on Nlg1. Whole-cell protein extracts were obtained by solubilizing cells in lysis buffer (50 mM HEPES, pH 7.2, 10 mM EDTA, 0.1% SDS, 1% NP-40, 0.5% DOC, 2 mM Na-Vanadate, 35 µM PAO, 48 mM Na-Pyrophosphate, 100 mM NaF, 30 mM phenyl-phosphate, 50 µM NH_4_-molybdate and 1 mM ZnCl_2_) containing protease Inhibitor Cocktail Set III, EDTA-Free (Calbiochem). Lysates were clarified by centrifugation at 8000 × g for 15 min. Equal amounts of protein (500 µg, estimated by Direct Detect assay, Merck Millipore) were incubated overnight with 2 µg rabbit anti-Nlg1 (Synaptic systems 129013), then precipitated with protein G beads (Dynabeads Protein G, Thermo Fisher Scientific) and washed 4 times with lysis buffer. At the end of the immunoprecipitation, 20 µL beads were resuspended in 20 µL of 2x loading buffer (120 mM Tris-HCl, 3% SDS, 10% glycerol, 2% β-mercaptoethanol, 0.02% bromophenol blue, pH = 6.8). After centrifugation, supernatants were separated on 4-15% Mini-PROTEAN TGX Precast Protein Gels (Bio-Rad) and transferred to nitrocellulose membranes for immunoblotting (semi-dry, 7 min, Bio-Rad). After blocking with 5% non-fat dried milk in Tris-buffered saline Tween-20 (TBST; 28 mM Tris, 137 mM NaCl, 0.05% Tween-20, pH 7.4) for 45 min at room temperature, membranes were probed for 1 h at room temperature or overnight at 4°C with mouse anti-phosphotyrosine (1:1000, Cell Signaling 9411S) or rabbit anti-Nlg1. After washing 3 times with TBST buffer, blots were incubated with horseradish peroxidase (HRP)–conjugated goat secondary antibodies (1:5000, Jackson Immunoresearch) or Easyblot (GeneTex) for IP for 1 h at room temperature. The latter was used to avoid the detection of primary antibodies from the IP. Target proteins were detected by chemiluminescence with Super signal West Femto (Pierce) on the ChemiDoc Touch system (Bio-Rad).

### Organotypic slice culture and single cell electroporation

Organotypic hippocampal slice cultures were prepared as described (Stoppini et al., 1991) from either wild type or Nlg1 knock-out mice (C57Bl6/J strain) obtained from N. Brose (MPI Goettingen). Animals were raised in our animal facility; they were handled and killed according to European ethical rules. Briefly, animals at postnatal day 5–8 were quickly decapitated and brains placed in ice-cold Gey’s balanced salt solution under sterile conditions. Hippocampi were dissected out and coronal slices (350 µm) were cut using a tissue chopper (McIlwain) and incubated at 35°C with serum-containing medium on Millicell culture inserts (CM, Millipore). The medium was replaced every 2-3 days. After 3–4 days in culture, slices were transferred to an artificial cerebrospinal fluid (ACSF) containing (in mM): 130 NaCl, 2.5 KCl, 2.2 CaCl_2_, 1.5 MgCl_2_, 10 D-glucose, 10 HEPES (pH 7.35, osmolarity adjusted to 300 mOsm). CA1 pyramidal cells were then processed for single cell electroporation using glass micropipets containing plasmids encoding TdTomato (6 ng/µl) and optoFGFR1 (13 ng/µl). For rescue experiments, a plasmid carrying the Nlg1 specific shRNA (16 ng/□l) was electroporated along with a resistant AP-Nlg1 (13 ng/µl), BirA^ER^ (ng/□l), TdTomato (6 ng/□l) and optoFGFR1 (13 ng/µl). Micropipets were pulled from 1 mm borosilicate capillaries (Harvard Apparatus) with a vertical puller (Narishige). Electroporation was performed by applying 4 square pulses of negative voltage (−2.5 V, 25 ms duration) at 1 Hz, then the pipet was gently removed. 10-20 neurons were electroporated per slice, and the slice was placed back in the incubator for 2-3 days before electrophysiology or confocal imaging.

### Immunocytochemistry

For visualization of recombinant AP-Nlg1 and spine morphology in electroporated CA1 neurons expressing tdTomato, AP-Nlg1 and BirA^ER^, organotypic slices were fixed with 4% paraformaldehyde-4% sucrose in PBS for 4 h before the permeabilization of membranes with 0.25% Triton in PBS. Slices were subsequently incubated with Atto647-conjugated Streptavidin (Molecular Probes, 1:200) for 2 hrs at room temperature. For visualization of HA-tagged optoFGFR1, fixed and permeabilized slices were incubated with rat anti-HA (Roche, clone 3F100, 1:100) overnight at 4°C. Slices were subsequently incubated with Alexa647-conjugated goat anti-rat antibody (Molecular Probes, 1:200) for 2 hrs at room temperature.

### Confocal microscopy and spine counting

For fixed slices, images of single CA1 electroporated neurons co-expressing tdTomato, BirA^ER^ and AP-Nlg1 (WT or Y782F mutant) were acquired on a commercial Leica DMI6000 TCS SP5 microscope using a 63x/1.4 NA oil objective and a pinhole opened to 1 time the Airy disk. Images of 4096 × 4096 pixels, giving a pixel size of 70 nm, were acquired at a scanning frequency of 400 Hz. The number of optical sections was between 150-200, using a vertical step size of 0.3-0.4 µm. The number of spines per unit dendrite length of tdTomato-positive cells in secondary and tertiary apical dendrites was calculated manually using Metamorph (Molecular Devices).

To assess the effect of optoFGFR1 stimulation on the formation of dendritic spines, we took confocal stacks of the dendritic tree of several CA1 neurons before light stimulation, then exposed the organotypic slices to dim 470 nm light pulses (1 s pulse every 1 s, 2.5 µW/mm^2^) through the LED array placed in the incubator for 24 h, and finally took another round of images of the same neurons. For such time-lapse imaging, short imaging sessions (10–15 min) of live electroporated slices were carried out with a commercial Leica DMI6000 TCS SP5 microscope using a 63x/0.9 NA dipping objective and a pinhole opened to one time the Airy disk. Slices were maintained in HEPES-based ACSF. Laser intensity in all these experiments was kept at a minimum and acquisition conditions remained similar between the two imaging sessions. 12-bit images of 1024 × 1024 pixels, giving a pixel size of 120 nm, were acquired at a scanning frequency of 400 Hz. The number of optical sections varied between 150-200, and the vertical step size was 0.3-0.4 µm. The number of spines per unit dendrite length of tdTomato-positive neurons was calculated manually in Metamorph.

### Electrophysiological recordings

Whole-cell patch-clamp recordings were carried out at room temperature in CA1 neurons from organotypic hippocampal cultures, placed on a Nikon Eclipse FN1 upright microscope equipped with a motorized stage and two manipulators (Scientifica). CA1 pyramidal neurons were imaged with DIC and electroporated neurons were identified by visualizing the GFP or Tdtomato fluorescence. The recording chamber was continuously perfused with ACSF bubbled with 95% O_2_ / 5% CO_2_ containing (in mM): 125 NaCl, 2.5 KCl, 26 NaHCO_3_, 1.25 NaH_2_PO_4_, 2 CaCl_2_, 1 MgCl_2_, and 25 glucose. 20 □M bicuculline and 100 nM NBQX were added to block inhibitory synaptic transmission and reduce epileptiform activity, respectively. We measured both AMPA-and NMDA-receptor mediated EPSCs upon electrical stimulation of Schaffer’s collaterals, using a double-patch clamp configuration to normalize the recordings with respect to a neighboring non-electroporated neuron(Shipman et al., 2011). Recordings were made under voltage clamp using the Multiclamp 700B amplifier (Axon Instruments). EPSCs and IPSCs were evoked in an electroporated neuron and a nearby non-electroporated neuron (control) using a bipolar electrode in borosilicate theta glass filled with ACSF and placed in the stratum radiatum or pyramidal layer; respectively. AMPAR-mediated currents were recorded at -70 mV and NMDAR-mediated currents were recorded at +40 mV and measured 50 ms after the stimulus, when AMPAR-mediated EPSCs are back to baseline. IPSCs were recorded at + 10 mV and in presence of 10 µM NBQX and 50 µM D-AP5 to block AMPARs and NMDARs, respectively. The series resistance Rs was left uncompensated, and recordings with Rs higher than 30 MΩ were discarded. EPSCs and IPSCs amplitude measurements were performed using Clampfit (Axon Instruments).

For LTP recordings, ACSF contained in (mM) 125 NaCl, 2.5 KCl, 26 NaHCO_3_, 1.25 NaH_2_PO_4_, 4 CaCl_2_, 4 MgCl_2_, 25 glucose, and 0.02 bicuculline, while recording pipettes were filled with intracellular solution containing in mM: 125 Cs-MeSO_4_, 10 CsCl, 10 HEPES, 2.5 MgCl_2_, 4 Na_2_ATP, 0.4 NaGTP and 10 phosphocreatine. Slices were maintained at 25°C throughout the recording. Baseline AMPAR-mediated EPSCs were recorded every 10 s for 2 min before LTP induction. Then LTP was induced by depolarizing the cells to 0 mV while stimulating the afferent Schaffer’s collaterals at 2 Hz for 100 s. Recordings were sampled every 10 s for 30 min after LTP induction.

### Computer simulations of AMPAR diffusion-trapping in LTP conditions

The computer program, written in C++, is based on a previous framework describing the role of AMPAR membrane dynamics in synaptic plasticity (Czöndör et al., 2012). Our original model included two types of processes to target AMPAR to synapses, i.e. diffusional trapping and vesicular recycling. However, based on recent experimental findings that hippocampal LTP primarily involves the diffusional trapping of extra-synaptic AMPARs (Granger et al., 2013; Penn et al., 2017), the current model focuses only on this process **(Fig. S5a)**. Briefly, a dendritic segment is approximated by a 2D rectangular region (2 µm × 10 µm) containing 5 synapses (squares of 0.3 µm × 0.3 µm, surface area ∼0.1 µm^2^), corresponding to a linear density of 0.5 synapse/µm as measured experimentally (Letellier et al., 2018). This area is populated with 1000 AMPARs, initially placed at random positions. AMPARs are characterized by their 2D coordinates x and y, over time, *t*. When AMPARs reach the region contours, rebound conditions are applied to keep them inside, i.e. the system is closed. At each time step (*Δt* = 100 ms), the coordinates are incremented by the distances Δx = (2D*Δt*)^1/2^ n_x_ and Δy = (2D*Δt*)^1/2^ n_y_, where n_x_ and n_y_ are random numbers generated from a normal distribution, and D is a diffusion coefficient which depends on whether AMPARs are outside (*D*_*out*_ = 0.1 µm^2^/s) or inside (*D*_*in*_ = 0.05 µm^2^/s) the synapse, values being taken from single molecule tracking experiments (Nair et al., 2013). Lower AMPAR diffusion within the synaptic cleft is attributed to steric hindrance. To introduce a diffusion barrier at the synapse (Ewers et al., 2014), AMPARs are allowed to cross the synaptic border with a probability P_crossing_ = 0.5.

Within the synapse, AMPARs may reversibly bind to static post-synaptic density (PSD) components, namely PDZ domain containing scaffolding proteins including PSD-95, S-SCAM, PICK or GRIP, through the C-terminal PDZ motifs of GluA1/2, or of TARPs (Bats et al., 2007; Kim and Sheng, 2004). To describe those dynamic interactions, we define two global parameters, the AMPAR/scaffold binding and unbinding rates (k_on_ = 1 s^-1^ and k_off_ = 0.02 s^-1^, respectively), obtained by previously fitting SPT and FRAP experiments (Czöndör et al., 2012). AMPARs are allowed to stay in the PSD if the probability of binding in this time interval (*k*_*on.*_*Δt*) is greater than a random number between 0 and 1 generated from a uniform distribution. Otherwise, AMPARs continue to diffuse with coefficient *D*_*in*_. When bound to the PSD, AMPARs move with a lower diffusion coefficient *D*_*trap*_ = 0.006 µm^2^/s, corresponding to confinement in the PSD (Czöndör et al., 2013; Nair et al., 2013). AMPARs stay in the PSD until their detachment probability (*k*_*off*_.*Δt*), exceeds another random number. Then, AMPARs can bind the same PSD again or escape into the extra-synaptic space **(Fig. S5b)**. At steady state (reached for *t* > 1/k_off_), there is a dynamic equilibrium between synaptic and extra-synaptic AMPARs **(Fig. S5c)**. The enrichment ratio between synaptic and extra-synaptic AMPAR density is given by the formula: P_crossing_ (D_out_ /D_in_) (1 + k_on_/k_off_). The maximal theoretical number of AMPARs per synapse is 200, when all extra-synaptic receptors in the system have been captured (given the excess of scaffolds versus AMPARs, we do not impose a saturation of binding sites here). With the chosen parameters however, there are about 30 AMPARs per synapse at basal state in control conditions, close to experimental measurements made by super-resolution imaging and freeze-fracture EM (Levet et al., 2015; Shinohara et al., 2008). The effect of Nlg1 tyrosine phosphorylation on basal synaptic AMPAR levels was simulated by raising the AMPAR/scaffold binding rate (k_on_), thereby mimicking an increase in the steady-state number of average post-synaptic AMPAR trapping slots observed experimentally (Giannone et al., 2013; Letellier et al., 2018; Mondin et al., 2011).

To simulate LTP, the AMPAR/scaffold unbinding rate (k_off_) was decreased at time zero from higher (0.02 to 0.08 s^-1^) to lower values (0.002-0.006 s^-1^), hereby mimicking a higher affinity of TARPs to PSD-95 induced by CamKII activation (Hafner et al., 2015; Opazo et al., 2010). When we tried instead to simulate LTP by raising the parameter k_on_ at time zero, the predicted time course was much more rapid than the one observed experimentally (i.e. the plateau was reached in about one minute). Thus, that type of mechanism is not likely to operate in the particular LTP protocol used here. The total length of the trajectories was set to 35 min, including a 5 min baseline, to match the whole duration of LTP experiments. Ten simulations were generated for each type of condition, and the number of AMPARs per synapse was determined and averaged (sem is within 1% of the mean). To determine the theoretical relationship between LTP level and basal synaptic AMPARs content, the parameter k_on_ was varied between 0.075 s^-1^ and 10 s^-1^, thus simulating synapses that contain less or more AMPARs, respectively.

### Sampling and statistics

For the analysis of dendritic spines observed by confocal microscopy, N et n values represent the total number of cells and dendrites, respectively. For each experiment, 3-4 independent dissections (from 2-3 animals) were used. Sample sizes were determined according to previous studies (Letellier et al., 2018; Shipman et al., 2011).

Summary statistics are presented as mean ± SEM (Standard Error of the Mean), including individual data points. Statistical significance tests were performed using GraphPad Prism software (San Diego, CA). Test for normality was performed with D’Agostino and Pearson omnibus normality test. Paired data obtained by imaging or electrophysiology experiments were compared using the Wilcoxon matched-pairs signed rank test when criteria for normality were not met. When paired data followed a normal distribution, we used a paired t-test. ANOVA test was used to compare means of several groups of normally distributes variables. Kruskal-Wallis test was used to compare several groups showing non-normal distributions. Dunn’s multiple comparisons post hoc test was then used to determine the p value between two conditions. Statistical significance was assumed when P<0.05. In the figures, *P<0.05, **P<0.01, ***P<0.001, ****P<0.0001.

## Ethical statement

The authors declare that they have complied with all relevant ethical regulations (study protocol approved by the Ethical Committee of Bordeaux CE50).

## Acknowledgements

We acknowledge L. Duschene, P. Scheiffele, and A. Ting for the generous gift of plasmids, N. Brose for the gift of Nlg1 KO mice, A. Hoagland for help with LED array construction and E. Isacoff for access to laboratory resources, M Sainlos for fruitful discussions, the Bordeaux Imaging Center (C. Poujol and S. Marais) for support in microscopy, the animal facility of the University of Bordeaux (in particular A. Lacquemant), the cell culture facility (especially S. Benquet, P. Durand and E. Verdier), and Biochemistry platform of the Neurocampus, J. Carrere and R. Sterling for technical assistance.

This work received funding from the Centre National de la Recherche Scientifique, Agence Nationale pour la Recherche (grant « Synthesyn » ANR-17-CE16-0028-01), Commission Franco-Américaine (Fulbright program), Conseil Régional Aquitaine (« SiMoDyn »), Investissements d’Avenir (Labex BRAIN), and Fondation pour la Recherche Médicale (« Equipe FRM » DEQ20160334916), and the national infrastructure France BioImaging (grant ANR-10INBS-04-01).

## Supplementary Figures

**Figure S1.**
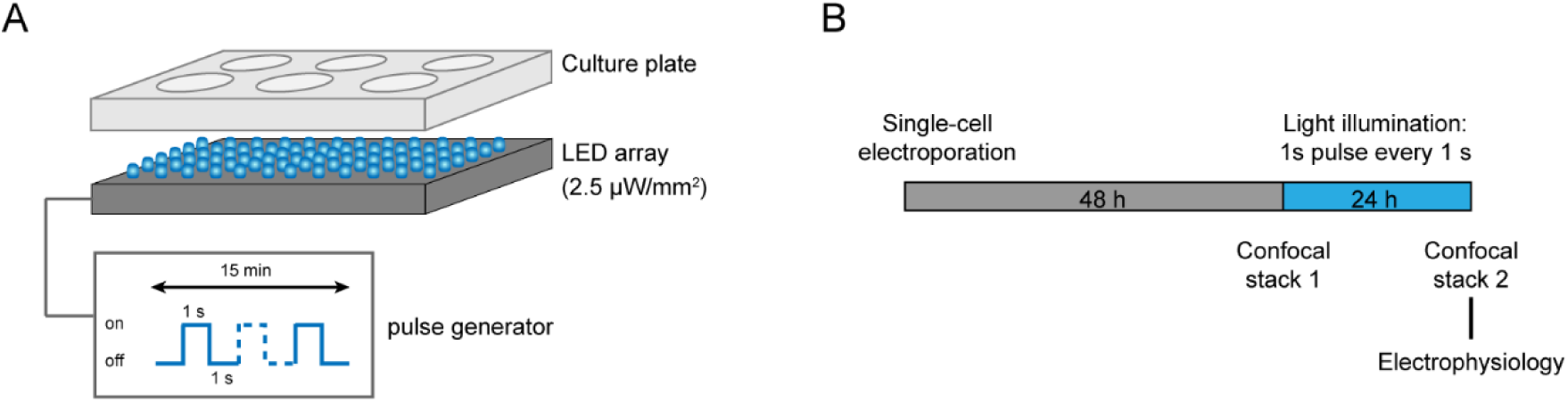
Experimental strategy for optogenetic stimulation. **(A)** Scheme representing the 470 nm light emitting diode (LED) array that is placed in a 37°C incubator and used to illuminate COS cells or organotypic slices contained in a 6-well plate. **(B)** Experimental procedure to investigate the effect of the optogenetic stimulation of optoFGFR1 on spine density and synaptic transmission. CA1 neurons in organotypic slices from WT or Nlg1 KO mice were electroporated at DIV 3-5 with tdTomato and HA-tagged optoFGFR1. Two days later, they were either stimulated with alternating blue light for 24 h or kept in the dark, and processed for imaging or electrophysiology.

**Figure S2.**
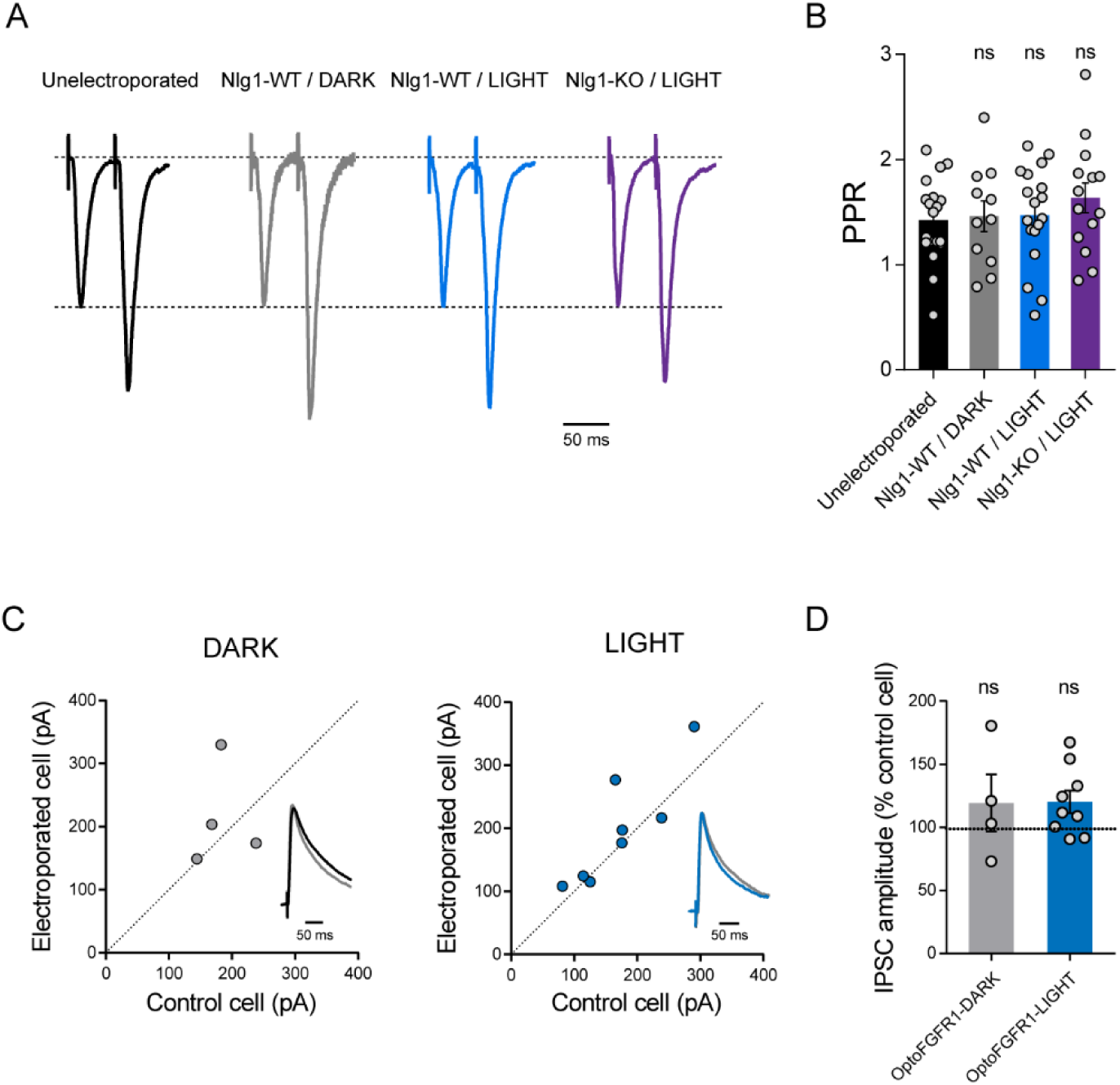
Stimulation of optoFGFR1 does not affect the paired pulse ratio (PPR) nor inhibitory currents in CA1 neurons. **(A)** Average EPSC traces in response to paired stimuli (50 ms interstimulus interval) recorded from an unelectroporated WT neuron (in black), WT neurons expressing optoFGFR1 kept in the dark (in grey) or exposed to light (in blue) and a Nlg1-KO neuron expressing optoFGFR1 and exposed to light (in violet). **(B)** Average PPR for the same conditions as in (A) (Wilcoxon matched-pairs signed rank test, ns: not significant). Data represent mean ± SE. **(C)** Scatter plots of IPSCs in optoFGFR1-expressing neurons versus paired unelectroporated neurons (control cell), in the light (blue) and dark (grey) conditions, respectively. Representative IPSCs traces (black or blue) were recorded at a holding potential of +10 mV and normalized to control (grey) and are shown as insets. **(D)** Average of IPSCs in the two conditions, normalized to the control (100%). Data were compared to the control condition by the Wilcoxon matched-pairs signed rank test. (ns, not significant).

**Figure S3.**
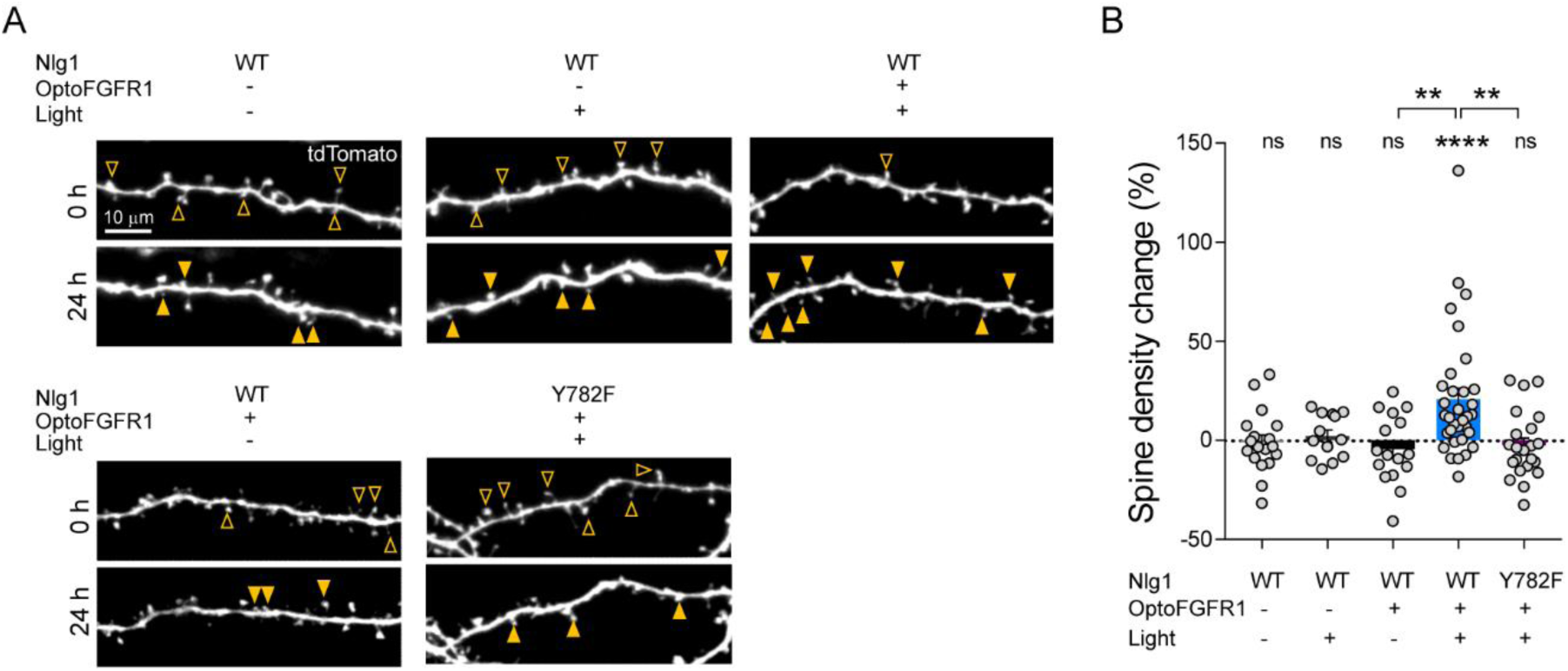
The light-induced increase in spine density is specific to Y782 phosphorylation in Nlg1. **(A)** Confocal images of tdTomato in apical dendrites from CA1 neurons before (0 h) and after (24 h) light activation. Control neurons either did not receive light, or did not contain optoFGFR1, or were expressing the Nlg1-Y782F mutant. Solid arrowheads point to spines which have appeared, and empty arrowheads to spines which have disappeared in the time period. **(B)** Normalized spine density for each condition (n = 13-33 dendrites from N = 4-10 cells). Change in spine density was assessed for each condition using the paired t-test (\*\*\*\**P* < 0.0001, ns: not significant). Spine density change was compared across conditions using the Kruskal-Wallis test followed by Dunn’s multiple comparison test, paired t-test (\*\**P* < 0.001).

**Figure S4.**
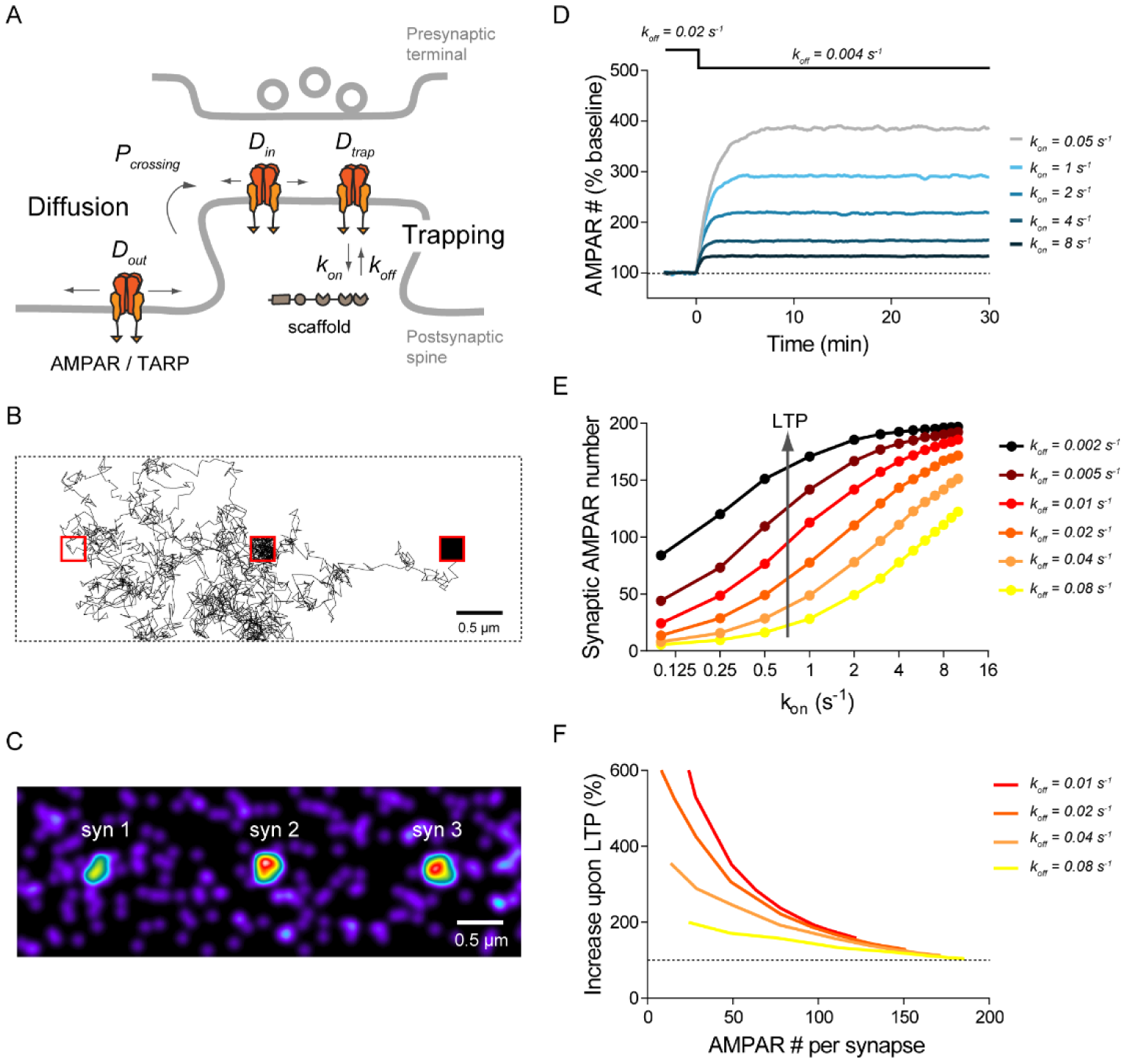
Computer simulations of AMPAR diffusion/trapping in LTP. **(A)** Schematic diagram of the AMPAR diffusion/trap process, with model parameters being indicated in italic. **(B)** Representative example of a single simulated AMPAR trajectory (time step *Δt* = 0.1 s, total duration 200 s). Note the fast AMPAR extra-synaptic diffusion and the slow diffusion when the AMPAR is trapped within the synapse (red squares). **(C)** Still image illustrating the steady-state distribution of AMPARs in the simulated neurite geometry. Individual AMPARs are represented by a Gaussian intensity profile of FWHM of 25 nm. Note AMPAR accumulation at synapses. **(D)** Graph showing the relative change in synaptic AMPAR content (in %) over time, when LTP is simulated by a drop in k_off_ at time zero. Curves correspond to different values of the final k_off_ parameter (initial k_off_ = 0.02 s^-1^). **(E)** Graph showing the steady-state synaptic AMPAR number versus the AMPAR/scaffold binding rate (k_on_ ranging from 0.5 s^-1^ to 10 s^-1^), for different values of the unbinding rate (k_off_ ranging from 0.005 s^-1^ to 0.2 s^-1^). The synaptic AMPAR level saturates at high k_on_ and/or low k_off_ values due to the depletion of the finite AMPAR extra-synaptic pool. **(F)** Predicted relationship between the basal synaptic AMPAR content and the LTP response, expressed as a percentage of the AMPAR baseline. The four curves represent different initial k_off_ values (the final value is 0.004 s^-1^ for all conditions).

